# optiPRM: A targeted immunopeptidomics LC-MS workflow with ultra-high sensitivity for the detection of mutation-derived tumor neoepitopes from limited input material

**DOI:** 10.1101/2023.08.22.554248

**Authors:** Mogjiborahman Salek, Jonas D. Förster, Jonas P. Becker, Marten Meyer, Pornpimol Charoentong, Yanhong Lyu, Katharina Lindner, Catharina Lotsch, Michael Volkmar, Frank Momburg, Isabel Poschke, Stefan Fröhling, Marc Schmitz, Rienk Offringa, Michael Platten, Dirk Jäger, Inka Zörnig, Angelika B. Riemer

**Affiliations:** Division of Immunotherapy and Immunoprevention, German Cancer Research Center (DKFZ) Heidelberg, Germany; Molecular Vaccine Design, German Center for Infection Research (DZIF), partner site Heidelberg, Germany; Faculty of Biosciences, Heidelberg University, Heidelberg, Germany; Antigen Presentation and T/NK Cell Activation Group, German Cancer Research Center (DKFZ) Heidelberg, Germany; Department of Medical Oncology, National Center for Tumor Diseases (NCT), NCT Heidelberg, a partnership between DKFZ and University Hospital Heidelberg, Germany; Clinical Cooperation Unit Applied Tumor Immunity, German Cancer Research Center (DKFZ) Heidelberg, Germany; Center for Quantitative Analysis of Molecular and Cellular Biosystems (Bioquant), Heidelberg University, Heidelberg, Germany; Immune Monitoring Unit, National Center for Tumor Diseases (NCT), NCT Heidelberg, a partnership between DKFZ and University Hospital Heidelberg, Germany; Clinical Cooperation Unit Neuroimmunology and Brain Tumor Immunology, German Cancer Research Center (DKFZ) Heidelberg, Germany; T Cell Discovery Platform, Helmholtz Institute for Translational Oncology (HI-TRON) Mainz – A Helmholtz Institute of the DKFZ, Mainz, Germany; Division of Translational Medical Oncology, German Cancer Research Center (DKFZ) Heidelberg, Germany; Division of Translational Medical Oncology, National Center for Tumor Diseases (NCT), NCT Heidelberg, a partnership between DKFZ and University Hospital Heidelberg, Germany; German Cancer Consortium (DKTK), DKFZ, core center Heidelberg, Germany; Institute of Immunology, Faculty of Medicine Carl Gustav Carus, TU Dresden, Dresden, Germany; National Center for Tumor Diseases (NCT), NCT Dresden, a partnership between DKFZ, University Hospital Carl Gustav Carus, Faculty of Medicine Carl Gustav Carus of TU Dresden and Helmholtz Center Dresden-Rossendorf, Dresden, Germany; German Cancer Consortium (DKTK), partner site Dresden, a partnership between DKFZ, University Hospital Carl Gustav Carus, Faculty of Medicine Carl Gustav Carus of TU Dresden, Helmholtz Center Dresden-Rossendorf and Max Planck Institute of Molecular Cell Biology and Genetics (MPI-CBG), Dresden, Germany; Division of Molecular Oncology of Gastrointestinal Tumors, German Cancer Research Center (DKFZ) Heidelberg, Germany; Department of General, Visceral and Transplantation Surgery, University Hospital Heidelberg, Germany; Department of Neurology, Medical Faculty Mannheim, Mannheim Center for Translational Neuroscience (MCTN), Heidelberg University, Mannheim, Germany; DKFZ Hector Cancer Institute at the University Medical Center Mannheim, Germany; Helmholtz Institute for Translational Oncology, Mainz (HI-TRON Mainz) – A Helmholtz Institute of the DKFZ, Mainz, Germany

**Author notes:** equally contributed (Mogjiborahman Salek, Jonas D. Förster).

## Abstract

Personalized cancer immunotherapies such as vaccines and T cell receptor (TCR)-transgenic T cells rely on the presentation of tumor-specific peptides by human leukocyte antigen (HLA) class I molecules to cytotoxic T cells. Such neoepitopes can for example arise from somatic mutations and their identification is crucial for the rational design of new therapeutic interventions. For their detection by liquid chromatography mass spectrometry (LC-MS), we have developed a parameter optimization workflow to tune targeted assays for maximum detection sensitivity on a per peptide basis, termed optiPRM. Optimization of collision energy using optiPRM allows for improved detection of low abundant peptides that are very hard to detect using standard parameters. Applying this to immunopeptidomics, we detected a neoepitope in a patient-derived xenograft (PDX) from as little as 2.5×10^6^ cells input. Application of the workflow on small patient tumor samples allowed for the detection of five mutation-derived neoepitopes in three patients. One neoepitope was confirmed to be recognized by patient T cells. In conclusion, we here present optiPRM, a targeted MS workflow reaching ultra-high sensitivity by per peptide parameter optimization, which allowed for the identification of actionable neoepitopes from sample sizes usually available in the clinic.

## BACKGROUND/INTRODUCTION

Personalized cancer immunotherapies such as vaccines and T cell receptor (TCR)-transgenic T cells rely on the presentation of tumor-specific peptides by human leukocyte antigen (HLA) class I molecules to cytotoxic T cells. Tumor-specific peptides, termed neoepitopes, can arise from somatic mutations or originate from non-canonical sources, *e.g.* translation of alternative open reading frames or non-coding regions of the genome, and their identification is crucial for the rational design of new therapeutic interventions ^1–4^.

Most neoepitope identification pipelines utilize next-generation sequencing data and subsequent candidate prioritization by *in silico* HLA binding prediction ^5^. Additionally, peptide binding to the respective HLA molecule and immunogenicity are often tested in *in vitro* assays. However, external loading as used in these *in vitro* approaches does not reflect the whole complexity of the antigen processing pathway including peptide generation by the proteasome, TAP transport into the endoplasmic reticulum (ER) and peptide loading to the HLA molecule. Therefore, these methods have only limited informative value as to whether a peptide is actually presented on the target cell. Currently, mass spectrometry (MS)-based immunopeptidomics is the only method to directly prove actual peptide presentation.

Although, the field of immunopeptidomics has made tremendous steps forward since it emerged about 30 years ago ^6–8^, many obstacles remain, both in terms of sample preparation and mass spectrometry analysis. For sample preparation, HLA class I-presented peptides are typically enriched either in a non-specific manner by mild acid elution (MAE) or HLA:peptide complexes are purified by immunoprecipitation (IP) before the presented peptides are separated from HLA molecules by acidic elution and purified by solid phase extraction (SPE), ultrafiltration and/or high-performance liquid chromatography (HPLC)^9^. While sample preparation by MAE is affected by the co-purification of contaminant cell surface peptides not associated with HLA class I molecules ^10^, sample preparation by IP suffers from losses of up to 99%, which make it necessary to use large amounts of input material to obtain the desired sensitivity for the detection of often low abundant neoepitopes ^11,12^. However, such large amounts are usually unavailable for patient samples, thus hampering the broader application of immunopeptidomics in clinical settings.

Subsequent MS analysis of isolated HLA peptides by untargeted methods, either by data-dependent acquisition (DDA) and more recently also data-independent acquisition (DIA) schemes, led to the unbiased identification of several thousands of HLA class I-presented peptides providing a global view of the immunopeptidome ^13–17^. However, these untargeted methods have limited sensitivity, leaving undetected both low-abundance peptides and those not well suited to ionization, including neoepitopes. In contrast, targeted approaches such as multiple reaction monitoring (MRM) and parallel reaction monitoring (PRM) can be utilized to detect HLA class I-presented peptides with much higher sensitivity compared to untargeted methods, but are limited to pre-defined peptide sets ^18–21^.

Here, we present optiPRM, a workflow for target-specific MS parameter optimization using direct infusion and apply it for systematic collision energy optimization. The accompanying data processing pipeline can be easily adapted for other MS parameters. Using our optiPRM assay tuned for maximum sensitivity and employing the per-target optimized parameters, we detect mutation-derived, immunogenic neoepitopes from limited input material of a patient-derived xenograft (PDX) cell line and sparse patient tumor samples with ultra-high sensitivity.

## MATERIALS & METHODS

### Cell lines

CaSki (ATCC CRL-1550) cells were obtained from ATCC and were cultivated in DMEM supplemented with 10% FBS, 1% penicillin/streptomycin and 2 mM L-Glutamine. SNU-17 cells were obtained from the Korean Cell Line Bank and were cultivated in RPMI-1640 medium supplemented with 10% FBS, 1% penicillin/streptomycin, 2 mM L-Glutamine and 1% HEPES. Cells were kept under standard conditions in a humidified incubator at 37°C and 5% CO_2_. For the immunoprecipitation of HLA:peptide complexes, cells were washed with ice-cold PBS and harvested using Accutase. Cells were frozen as dry pellets in liquid nitrogen until further usage.

### Patient-derived xenograft cell line

Generation of the xenograft and the derived cell line TIPC309 was performed as described previously ^22^. As this is not a commercially available cell line, it has not undergone a formal authentication process. The descent of a cell line from a primary tumor was verified by tracing mutations detected in primary tumor/xenograft by PCR and Sanger sequencing. For the +IFNγ experiments, the cell line was treated with 333 IU/ml IFNγ for 48 h before harvest. Cells were washed twice with PBS supplemented with 1% BSA, and the dry cell pellet was snap frozen in liquid nitrogen.

### Patient material

For this study, patient tumor material was collected from 5 patients. All patients provided informed consent approved by ethics votes (S-206/2011 (MASTER trial) and S-205/2007 (NCT Biobank), Ethics Committee of the Medical Faculty of Heidelberg University; 2019-643N (BrainTUNE), Ethics Committee of the Medical Faculty Mannheim of Heidelberg University). Patient information is summarized in Supplementary Table S1.

### Immunoprecipitation of HLA:peptide complexes

Immunoprecipitation (IP) of HLA:peptide complexes was based on previously published protocols ^15,23^ and the optimization reported in Figure 1. Cell line/PDX samples were lysed with a ratio of 1 ml lysis buffer per 1×10^8^ cells, composed of 1% N-octyl-β-D glucopyranoside, 0.25% sodium deoxycholate, protease inhibitor cocktail (Sigma-Aldrich, St. Louis, MI, USA)/phenylmethylsulfonyl fluoride in PBS. Snap-frozen tissue samples were homogenized in 1 ml lysis buffer per 100 mg tissue on ice in 3 – 5 short intervals of 5 s each using an Ultra Turrax homogenizer (IKA, T10 standard, Staufen, Germany) at maximum speed, as previously described ^24^. After centrifugation at 40,000×g at 4°C for 30 min, HLA:peptide complexes were immunoprecipitated by incubation with mouse anti-human HLA-A, -B, -C monoclonal antibody (mAb)(clone W6/32; Biolegend, San Diego, CA, USA) crosslinked to protein A Beads (Protein A Sepharose 4B, Invitrogen, Waltham, MA, USA) or, where indicated, protein G Beads (GammaBind Plus Sepharose beads, Cytiva, Marlborough, MA, USA) for 4 h, at 4°C under constant mixing on a rotating wheel. The mAb/beads and mAb-beads/cells ratios were 125 µg/50 µL and 170 µL 50:50 beads suspension per 1×10^8^ million cells, respectively. Supernatant was discarded after centrifugation at 3200×g, for 3 min at RT. Pelleted mAb-beads with bound HLA:peptide complexes were washed 3X with ice-cold 20 mM Tris-HCl (pH 8) containing 150 mM NaCl, then 3X with ice-cold 20 mM Tris-HCl (pH 8) containing 400 mM NaCl, and finally 3X with 20 mM Tris-HCl (pH 8) alone. Immunoprecipitation success was evaluated by comparing HLA class I abundance in input and flowthrough samples by SDS-PAGE with subsequent Western Blot analysis using a pan-HLA class I antibody (clone EP1395Y; GeneTex, Irvine, CA, USA).

**Figure 1:**
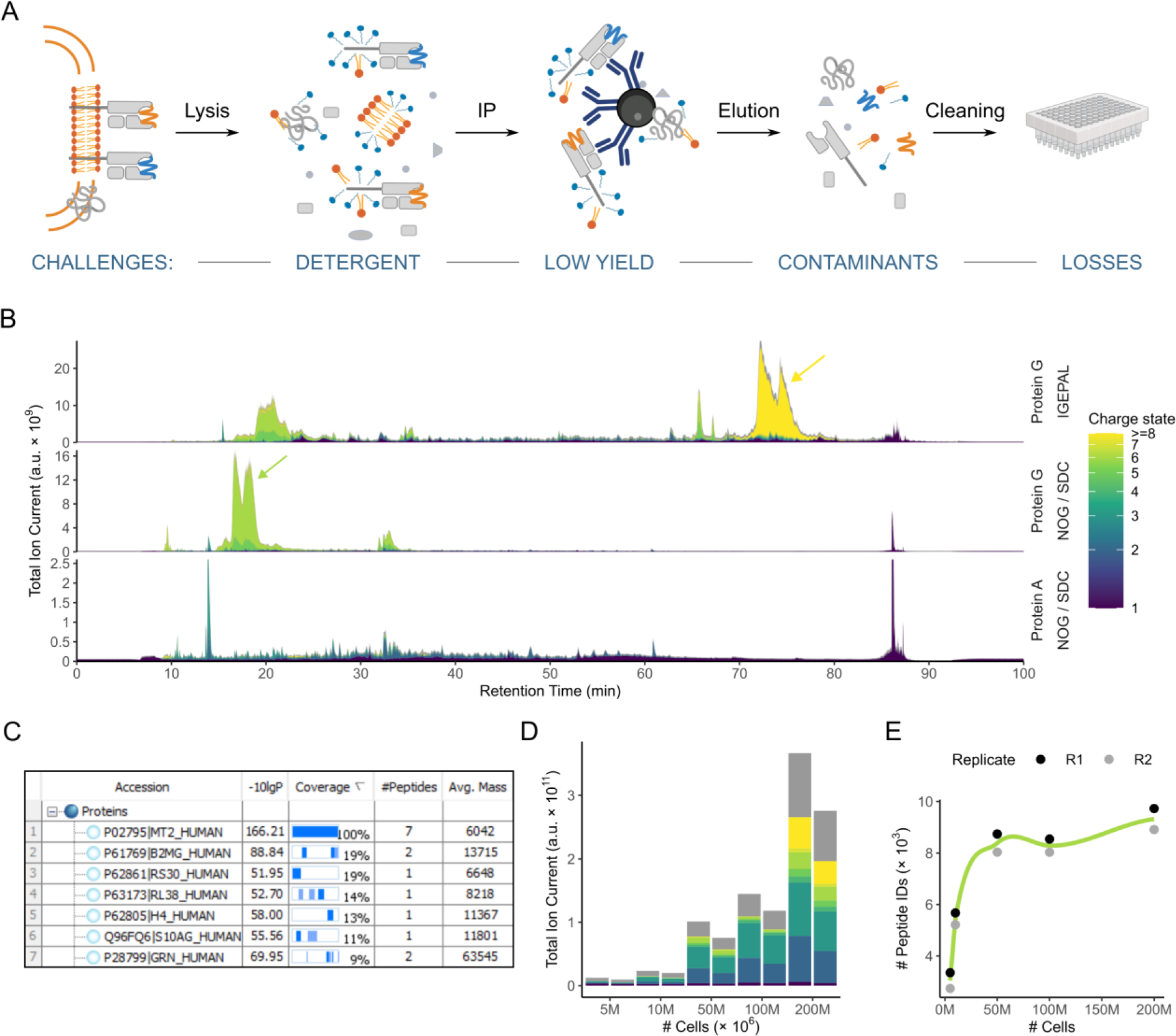
Optimization of HLA:peptide extraction and sample preparation workflow. (A) Schematic overview of key steps of the workflow and associated challenges. (B) Total ion chromatograms resulting from indicated immunoprecipitation (IP) conditions (detergents: polydisperse: IGEPAL CA-630, non-polymeric: n-octyl-β-D glucopyranoside/sodium deoxycholate (NOG/SDC), IP beads: protein A or protein G). The color code indicates charge states from +1 to higher than or equal to +8 (unknown charge states are shown in grey). The yellow arrow highlights contamination with high charge states (z ≥ +8), and the green arrow is a separate contamination with a majority charge state z = 6. (C) Protein detections resulting from DDA acquisition of an IP (NOG/SDC, protein G) after tryptic digestion. (D) Quantification of the total signal at indicated charge states (color-coded as in (B)) resulting from NOG/SDC protein G IPs with the increasing number of cell load. (E) The number of unique peptides identified in (D) as a function of the increasing number of cells.

### Peptide purification

HLA:peptide complexes were eluted from the mAB-beads with 1 ml 0.3% trifluoroacetic acid (TFA) and desalted by solid-phase extraction using a 100 mg SepPak tC18 96-well plate (Waters, Milford, MA, USA). An optional oxidation step was applied where indicated while the sample was bound to the SepPak cartridge by a brief application of 0.45% performic acid. After elution with 28% acetonitrile (ACN), samples were dried by vacuum centrifugation. Specific conditions are listed in Supplementary Table S2. For indicated samples, peptides were purified using restricted access media (RAM; MAYI-ODS 50μm, 30×4, 6 mm; Shimadzu, Kyoto, Japan) as described before with minor modifications ^25^. The protocol was tested in acidic and neutral conditions and with or without drying by vacuum centrifugation. Ordinary 200 µL pipette tips were prepared as described elsewhere ^26^ with a piece of glass fiber filter (Macherey-Nagel, Düren, Germany) acting as a frit wedged deep into the tip. The pipette tip was then filled with 7 mg of RAM resuspended in 10 mM ammonium acetate (AA) containing 60% ACN. The liquid was removed to compact the RAM material in a solid phase by pushing with a Luer lock syringe fitted on the top of the pipette tip. In acidic conditions, the 10 mM ammonium acetate was replaced by 0.1% TFA. A second wash was carried out with 200 μL 10 mM ammonium acetate (or 0.1% TFA in acidic conditions) in 5% ACN. Dried IP eluates were solubilized in 10 mM ammonium acetate (or 0.1% TFA) containing 5% ACN and passed through the solid phase. The solid phase was washed twice with 150 μL 10 mM ammonium acetate (or 0.1% TFA) in 5% ACN. Peptides were eluted in 50 μL 10 mM AA (or 0.1% TFA) in 60% ACN. In conditions where the IP eluate was not dried by vacuum centrifugation before purification, the acidic pH of the eluate was neutralized with 20 µl 1 M AA.

### Whole exome sequencing

The DNA libraries of the tumor and matched control samples were sequenced on NovaSeq 6000 (2x 100 bp), demultiplexing of the sequencing reads was performed with Illumina bcl2fastq (2.20). Adapters were trimmed with Skewer (version 0.2.2)^27^. Sequencing reads were aligned using the DKFZ alignment workflow from the ICGC Pan-Cancer Analysis of Whole Genome projects (DKFZ AlignmentAndQCWorkflows v1.2.73, https://github.com/DKFZ-ODCF/AlignmentAndQCWorkflows). The human reference genome version GRCh37/hg19 was used.

### RNA sequencing and expression quantification

RNA libraries of tumor samples were prepared using Kapa RNA HyperPrep Kit with RiboErase (Roche, Basel, Switzerland). Samples were sequenced on NovaSeq 6000. RNA-seq reads were aligned and gene expression quantified using the DKFZ RNAseq workflow (v1.2.22-6, https://github.com/DKFZ-ODCF/RNAseqWorkflow) as previously described ^28^. For total library abundance calculations, during TPM and FPKM expression values estimation, genes on chromosomes X, Y, MT, and rRNA and tRNA were omitted as they can introduce library size estimation biases ^28,29^.

### Mutation calling and annotation

Somatic small variants (SNVs and InDels) in matched tumor-normal pairs were called using the DKFZ in-house pipelines (SNVCallingWorkflow v1.2.166-1, https://github.com/DKFZ-ODCF/SNVCallingWorkflow; IndelCallingWorkflow v1.2.177, https://github.com/DKFZ-ODCF/IndelCallingWorkflow) as previously described ^30^. Raw calls for InDels were obtained from Platypus ^31^. The proteins coding effect of SNVs and InDels from all samples were annotated using ANNOVAR ^32^ according to GENCODE gene annotation (version 19)^33^ and overlapped with variants from dbSNP10 (build 141) and the 1000 Genomes Project database. Mutations of interest were defined as somatic SNV and InDels that were predicted to cause protein coding changes (non-synonymous SNVs, gain or loss of stop codons, splice site mutations, frameshift and non-frameshift InDels)^28^. Arriba ^34^ was used to detect gene fusions from RNA-Seq data.

### Epitope prediction and prioritization

After mutation calling, mutated protein sequences were generated (21mers with mutated amino acid in the middle for SNVs, frameshift sequences with 10 wildtype amino acids upstream of the mutation site for InDels and fusions) and netMHCpan 4.1 predictions ^35^ were performed for 8-11mers for the HLA alleles of the corresponding sample. Of these, 50 – 70 candidates were selected based on criteria such as eluted ligand rank and binding affinity, expression level and mutated allele frequency based on RNA-seq, as well as position and nature of the mutated amino acid(s) with a preference for peptides presented by HLA-A and -B.

### Synthetic peptides

Custom-synthetized stable isotope labelled (SIL) peptides were designed so that each contains at least one heavy amino acid and so that no peptide falls into the same precursor isolation window (ca. ± 1 m/z) as another nor contains the sequence of another peptide in full, in an unlabeled manner. Peptides were obtained from JPT Peptide Technologies (Berlin, Germany) or Synpeptide Co., Ltd (Shanghai, China) with low chemical purity (> 70%). The isotope purity is higher than 99 atom% ^13^C and ^15^N, and incorporated heavy amino acids are V (^13^C_5_, ^15^N_1_), L/I (^13^C_6_, ^15^N_1_), A (^13^C_3_, ^15^N_1_), F (^13^C_9_, ^15^N_1_), K (^13^C_6_, ^15^N_2_) and R (^13^C_6_, ^15^N_4_).

### Optimization of detection parameters by direct infusion MS

The set of predicted target synthetic peptides was first analyzed by direct infusion MS (DI-MS) in order to optimize normalized collision energy (NCE) for each precursor in a mixture of peptides. For that, dried synthetic peptides were solubilized in 50% ACN / 4% formic acid (FA) to a final concentration of 1 pmol/μL. 1 – 5 μL of the sample was drawn into a fused silica capillary with the help of a syringe (Hamilton, Bonaduz, Switzerland). The sample was then channeled into the emitter through a liquid junction connected to the fused silica capillary mounted in front of the MS instrument (patent application WO2023104771). The spray voltage was set to 1100 V and ramped up to where a stable spray was achieved. NCE values ranged from 4% to 42% and each value, in 2% steps, was measured 4 times for four charge states (1+ to 4+) of each peptide. Orbitrap Exploris 480 (Thermo Fisher Scientific, Waltham, MA, USA) in targeted MS2 scan was operated using a resolution of 30.000 at 200 m/z, standard automated gain control (AGC) target, 350 ms injection time (IT) in centroid data acquisition mode. For all measurements, the isolation window was 0.4, 0.7, 1 or 1.5 m/z, depending on the precursor mass, which are the smallest windows the manufacturer recommends. Ion traces of all expected transitions were extracted with 12 ppm mass tolerance from the centroided spectra and deconvoluted using purpose-made R scripts (https://github.com/jonasfoe/ms_targeted_workflows). The best NCE value was selected so that the intensity of any 5th most intense transitions was maximized, thereby maximizing the chance of detecting at least 5 transitions.

### LC-MS

All samples were analyzed by liquid chromatography (U-3000, Thermo Fisher Scientific) coupled to Orbitrap Exploris 480 (Thermo Fisher Scientific) using a nanoEase M/Z Peptide BEH C18 Column, 130Å, 1.7 µm, 75 µm X 200 mm LC column (Waters, Milford, MA, USA). For dilution experiments, samples were measured with increasing input to avoid carryover.

For reference acquisition, peptides were dissolved in 5% ACN, 0.1% TFA and injected at 2 – 200 fmol per peptide on column. First, the retention time (RT) for each target peptide was determined by LC-MS analysis of synthetic SIL peptide mixtures in DDA mode with an inclusion list of all target peptides in all four charge-states. The resolution was set to 120,000 at 200 m/z, 3×10^6^ AGC target, 50 ms maximum IT. The measured RT times were tested for the expected linear variation versus the hydrophobicity index (HI). Therefore, sequence-specific HIs were calculated for all target peptides with the SSRCalc tool ^36^. Only unmodified peptides were included and parameters were set to 100 Å C18 column, 0.1% formic acid separation system, and without cysteine.

IP samples were dissolved in 2.5 – 5 µl of 5% ACN in 0.1% TFA with 3min batch sonication and spiked with 50 – 125 fmol Peptide Retention Time Calibration (PRTC) Mixture (Pierce Biotechnology, Rockford, IL, USA). To curb surface losses, 375 fmol BSA Protein Digest BSA digest (Pierce Biotechnology) were added or QuanRecovery Vials with MaxPeak HPS (Waters) were used. The described BSA matrix (free of target peptides) was systematically injected before IP samples as a negative control. LC solvents were Solvent A (0.1% FA in H_2_O) and Solvent B (100% or 80% ACN, 0.1% FA) with specific LC gradients listed in Supplementary Table S2.

MS data was acquired with a resolution of 60,000 at 200 m/z, over the mass range from 150 to 1450 m/z, with 3×10^6^ AGC target and 25 ms maximum IT. MS2 data was acquired with PRM scans using a resolution of 60,000 at 200 m/z. The target precursor list was provided with preselected charge states, the corresponding m/z and optimized collision energy values for each target, and their expected retention time (+/− 1.5 min) pre-defined with SIL peptides and indexed to PRTC peptides. The normalized AGC target was set to 1000% (or 1×10^6^). The maximum injection time mode was set to dynamic, aiming for a coverage of at least 5 points across the chromatographic peak. The dynamic RT feature using the PRTC mixture was active. Protonated polycyclodimethylsiloxane (PCM-6, a background ion originating from ambient air) at 445.12 m/z served as a lock mass. MS2 data was acquired with PRM scans using a resolution of 60,000 at 200 m/z with a narrow isolation window (≤ 1 m/z) tuned per precursor. Heavy precursor settings were 30 ms IT, 5 × 10^5^ AGC target. Light targets were measured with 1000 ms IT, 1 × 10^6^ AGC target. These settings are identical to those we used to analyze endogenous peptides in a biological sample.

### Data Analysis

Targeted LC-MS data was analyzed using Skyline software (v. 20.2)^37^. Transitions were extracted in centroided mode with 7 ppm mass tolerance. Detected peaks were manually curated. Light peaks were discarded when peak retention times or shapes did not match the heavy reference, when the normalized spectral contrast angle (NSA)^38^ was low, or when too few transitions were detected. The top 12 transitions of the spectral library were considered for evidence scoring. Transitions were deselected if they were clearly perturbed from a closely coeluting peak. If the peak intensity did not allow for the detection of all transitions, NSA calculation was limited to a minimum of the top 5 transitions. Data evaluation for parameter optimization and detection diagnostics was performed with R (v. 4.2)^39^ with the “tidyverse” suite of packages (v. 2.0.0)^40^. The scripts are published alongside this publication on GitHub at https://github.com/jonasfoe/ms_targeted_workflows.

Untargeted experiments were analyzed via database search using PEAKS X Pro (Bioinformatics solutions Inc., Waterloo, ON, Canada) using unspecific or tryptic digest mode, variable modifications Oxidation (M) and Acetylation (N-term) and the UniProt human reference proteome (2020-04-22).

The mass spectrometry proteomics data have been deposited to the ProteomeXchange Consortium via the PRIDE ^41^ partner repository with the dataset identifier PXD043542 and DOI: 10.6019/PXD043542.

### Recombinant soluble HLA:peptide complex production

Soluble HLA:peptide complexes were generated by assembling disulfide-trapped single-chain trimers (SCT)^42^, where neoepitopes identified by optiPRM were fused in sequence to the human beta-2-microglobulin (β2M) domain and the predicted binding HLA ectodomain, each domain being linked with various G_X_S linker motifs as previously described in ^43^ with modifications. In all constructs, the sequence coding for the linker TSTGQL***HHHHHHHH***QL**GLNDIFEAQKIEWHE***LVPRSLVPRS*TS, including a His8-tag (**bold**, *italics*), a BirA biotin ligase recognition site (**bold**) and a double thrombin protease cleavage site (*italics*), was inserted between the HLA class I ectodomain and the Fc portion of mouse IgG2a. For protein production, plasmids encoding for various SCTs were co-transfected with a plasmid encoding for an ER-retained BirA-ligase into FreeStyle 293-F cells (Invitrogen) using the 293-free transfection reagent (Merck, Darmstadt, Germany) according to the manufactures’ protocols. Transfected cells were maintained overnight in FreeStyle 293 expression medium (Invitrogen) supplemented with 4 µg/ml D-biotin (Sigma-Aldrich) at 37°C, 8% CO_2_ and 100 rpm with a 50 mm shaking diameter. The next day freshly dissolved valproic acid (VPA)(Sigma-Aldrich) was added to the transfected culture to a final concentration of 4 mM as well as antibiotic-antimycotic solution (Sigma-Aldrich). The supplemented culture was further maintained for 6 days before the harvest of the cell supernatant. Cell-free supernatant was supplemented with 0.1 volumes of 10X Dulbecco’s PBS (Sigma-Aldrich) and 2 units thrombin (Merck)/mg SCT protein previously measured by mouse-IgG-Fc-based sandwich ELISA following an overnight incubation at 37°C. Soluble monomeric SCT molecules were further purified by immobilized metal affinity chromatography (IMAC) using Ni-INDIGO MagBeads (Cube Biotech, Monheim, Germany) according to the manufacturer’s instructions. Eluted proteins were finally dialyzed against PBS (pH 7.4) and their purity and metabolic biotinylation were verified by a non-reducing 10% SDS-PAGE in the presence and absence of streptavidin. In SCT 13806, a sequence coding for the peptide RRIRASQLLLH (Fusion26[breakpoint + amino acids 19–29]), in SCTs 13808 and 13809, a sequence coding for the peptide ARFMSPMVF (Fusion24[breakpoint + amino acids 2–10]), and in SCT 14224, the sequence coding for the peptide APRQPLSSI (RNF111[A609–I617/S611R]) were cloned, respectively. In SCTs 13806 and 13808, the leader-less ectodomain HLA-B*27:05[Y84C], in SCT 13809, the leader-less ectodomain HLA-C*07:02[Y84C], and in SCT 14224, the leader-less ectodomain HLA-B*56:01[Y84C] were cloned, respectively.

### Patient sample neoantigen-specific T cell identification

Monomeric biotinylated SCTs were mixed at a 4:1 ratio with different streptavidin (SA)-fluorochrome conjugates comprising SA-R-phycoerythrin (SA-RPE, Miltenyi, Bergisch Gladbach, Germany), SA-allophycocyanin (SA-APC, BioLegend), SA-Brilliant Violet 421 (SA-BV421, BD Biosciences, Franklin Lakes, NJ, USA) and SA-Brilliant Ultra Violet 395 (SA-BUV395, BD Biosciences) to form pMHC-I multimers. To increase the staining specificity and to allow multiplexed analysis of various epitopes in one staining, individual SCTs were complexed with two different SA-fluorochrome conjugates representing a unique dual-color combination as described before ^44^. For the *ex vivo* detection of neoantigen-specific T cell populations, cryopreserved patient PBMCs were thawed and rested overnight. CD8^+^ T cells were enriched using the REAlease® CD8 MicroBead Kit (Miltenyi) according to the manufacturer’s recommended protocol. Alternatively, CD14^+^ and CD25^+^-depleted PBMC were cultured in AIM-V media (Thermo Fisher Scientific) supplemented with human serum in the presence of FLT3L (Miltenyi), 9-10mer neoantigen peptides (2 µM, DKFZ peptide synthesis facility), TNF-α (PeproTech, Cranbury, NJ, USA), IL-1β (PeproTech), PGE1 (Sigma-Aldrich), IL-7 (Miltenyi) and IL-15 (Miltenyi) for 10-14 days as described previously ^21^. *Ex vivo* CD8^+^ T cells or *in vitro* stimulated (IVS) cultures were labeled with the Zombie Aqua™ Fixable Viability Kit (BioLegend) to exclude dead cells in all flow cytometry experiments. Next, Human TruStain FcX (Fc receptor blocking solution, BioLegend) was used to avoid nonspecific binding. Cells were then stained with the pMHC-I multimer libraries at room temperature for 25 min. After one wash, cells were stained by a cocktail containing optimal titrated antibodies (all from BioLegend) against human CD14 (M5E2), CD16 (3G8), CD19 (H1B19), and CD335 (9E2)(Brilliant Violet 510 for all); CD8 (SK1) APC-Cy7 and CD3 (UCHT-1) Alexa Fluor 700. Finally, the stained cells were stored in DPBS supplemented with 2.5% (v/v) paraformaldehyde (PFA) and 1% FCS before flow cytometry measurement on a BD LSRFortessa flow cytometer (BD Biosciences). pMHC-I multimer binding T cells were identified by a Boolean gating strategy in FlowJo (BD Biosciences) v.10.8.1 software as live CD8^+^ T cells stained positively in two pMHC multimer channels and negatively in all other pMHC multimer color channels, as previously described ^45^.

### 3D structural modelling of HLA:peptide complexes

3D structural models for binding of epitopes APRQLPSSI and APSQLPSSI to HLA-B*56:01 were generated using PANDORA 2.0.0b2 ^46^ with template 4U1K, loop_models 100, loop_refinement very_slow and restraints_stdev 0.3 and the best model by molpdf score was used.

## RESULTS

### Optimization of key and alternative parameters during sample preparation

Sample preparation in immunopeptidomics, *i.e.* the purification of HLA class I-presented peptides, has been described as the “Achilles’ heel” of the whole workflow ^47^ posing several challenges (Fig. 1A). First, the choice of detergent impacts the efficiency of the cell lysis and thus the yield of capturing HLA:peptide complexes ^11^. Additionally, inappropriate choice of detergent and its concentration can cause contamination, clogging of LC columns and reduced electrospray ionization ^48^. Second, the choice of resin for antibody coupling affects the amount and nature of unwanted background signals. Third, sample clean-up by conventional SPE with C_18_ material and unspecific elution by organic solvent leads to losses, particularly of highly hydrophilic peptides which are not retained and highly hydrophobic peptides which cannot be separated from the proteinaceous background.

We addressed the aforementioned challenges by revisiting and comparing previously published immunopeptidomics sample preparation protocols. To this end, we first tested six detergents in three concentrations (0.1 – 1%) for their ability to release HLA class I molecules from CaSki cell lysates. The detergents chosen were all mild and non-denaturing, *i.e.* not breaking protein-protein interactions and have been previously reported for the purification of membrane proteins ^10^. According to our data, higher detergent concentration led to a higher HLA class I signal as determined by Western blot analysis with the highest signals obtained for lysis by 1% n-octyl-β-D glucopyranoside (NOG) with 0.25% sodium deoxycholate (SDC), a combination previously published for the identification of HLA class I-presented peptides including neoepitopes ^13^, followed by lysis with 1% 3-[(3-cholamidopropyl)dimethylammonio]-1-propanesulfonate (CHAPS)(Supplementary Fig. S1A). Further analysis of two technical replicates of these samples by DDA MS revealed a mean of 5581 unique peptides for the samples lysed by CHAPS compared to 8008 unique peptides for the samples lysed by NOG/SDC (Fig. S1B). In addition, we assessed the optimal beads:antibody ratio by incubating varying amounts of panHLA antibody W6/32 (0 – 150 µg) with 25 µg of either Protein A or Protein G dry beads. Using oriole staining and quantification of bound antibody on beads and unbound fraction in the supernatant (Fig. S1C), we found 125 µg antibody per 25 µg of dry beads to be an optimal compromise, which also agrees with previously published protocols ^49^. Notably, the optimal antibody amount determined here is much lower than the amount suggested by the manufacturer.

To allow for an effective assessment of peptide purification performance from the LC-MS data, we color-coded the total ion current to highlight ion signals that diverge from the typically expected charge range of +1 – +3 for short HLA class I peptides. Notably, samples lysed using the polydisperse detergent octylphenoxypolyethoxyethanol (IGEPAL/Nonident P-40), previously used for the extraction of HLA class I-presented peptides by us and others ^20,50^, showed contamination by highly charged proteinaceous background (here mainly consisting of beta-2-microglobulin (F5H6I0)) and frequently caused clogging of LC columns (Fig. 1B, top panel, yellow arrow). Using NOG/SDC in the lysis buffer, the contaminations were limited to an early eluting species of ions with charge state ≥ +4 at ca. 6 kDa (Fig. 1B, middle panel, green arrow) and LC clogging was completely resolved. Trypsin digest of a sample and subsequent MS analysis indicated that the early eluting background signals originate from Metallothionein-2 (P02795, Fig. 1C), which has a hydrophobicity in the range expected for HLA class I-presented peptides and is therefore not removed by SPE using conventional C_18_ material. This contamination was resolved when performing the assay with Protein A beads instead of Protein G beads (Fig. 1B, bottom panel).

Contaminating proteins, which typically appear at late retention time and high charge states (e.g. Fig. 1B, yellow arrow) should in principle be retained on the SPE C_18_ material, when eluting peptides with a limited ACN concentration of 28%. However, when we tested the capacity of 100 mg sorbent SepPak cartridges with lysates from 5 – 200×10^6^ cells, we found an increasing amount of highly charged precursor ions with 200×10^6^ CaSki cells as input material, suggesting a diminished capability of the cartridge to retain protein contamination when overloaded (Fig. 1D). DDA analysis of these samples showed an increase in peptide identifications with increasing amount of input material which is plateauing after 50×10^6^ cells input material (Fig. 1E). Therefore, subsequent experiments were performed with Protein A beads and NOG/SDC in the lysis buffer. To avert overloading of SepPak cartridges, the amount of input material was adjusted accordingly.

### Systematic optimization of normalized collision energy improves the sensitivity of peptide detection

DDA MS is commonly used for the detection of HLA class I-presented peptides. However, DDA has limited depth of analysis and typically peptides at the limit of detection (LOD) are identified inconsistently or not at all. PRM alleviates this limitation by targeted acquisition of peptides of interest (Fig. 2A)^51^. To further increase the sensitivity of PRM, we implemented a pipeline for the systematic optimization of various instrument parameters utilizing direct infusion (DI-)MS. In DI-MS, peptides are analyzed without LC separation, thus allowing for systematic scanning of virtually all available values for a given parameter and peptide. DI-MS is usually performed with a mixture of target peptides (>30) allowing for rapid iteration through precursors. We used this pipeline for the optimization of normalized collision energy (NCE). The formula for NCE incorporates precursor charge state and mass information to adjust the collision energy in a generalized fashion. However, we found that the resulting energy is not generally optimal, especially for the typically non-tryptic peptides presented by HLA class I molecules. We scanned the NCE range from 4 – 42% using DI-MS to find an optimal NCE for a given precursor. To maximize the chance of detecting at least 5 transitions per peptide, the optimal NCE was set to maximize the intensity at a depth of 5 transitions into the spectrum. Transitions smaller than 3 amino acids were excluded as these ions have limited sequence information and their MS signals are susceptible to background chemical noise. To illustrate NCE optimization using DI-MS, the data for the non-tryptic peptide DLQPETTDL[+7]Y^++^ is presented (Fig. 2B). NCE of 14% — compared to the commonly used NCE of 30% — yielded much higher MS2 signal intensity. The MS2 spectrum recorded for this peptide during LC-MS analysis verified that fragmentation at the optimum NCE yields much higher sequence coverage due to the generation of larger fragment ions, while the spectrum generated at 30% NCE is dominated by rather small fragment ions (Fig. 2C). The corresponding extracted ion chromatograms (XICs) of the representative peptide spiked into a HeLa digest at the limit of detection further illustrates the benefit of the optimized NCE (Fig. 2D). To quantify the gain in sensitivity, we performed a dilution series down to 1.4 amol of synthetic peptide in a matrix of consistent amount of HeLa digest. We evaluated peptide detection based on the normalized spectral contrast angle (NSA)^38^ of the top 5 reference transitions. Here, an approximately 12 times higher concentration was needed when using an NCE of 30% to reach an NSA ≥ 0.85 for confident identification when compared to the optimized NCE of 14% (Fig. 2E). To assess the benefit of NCE optimization using DI-MS on signal intensity for a total of 1343 putative HLA class I-presented synthetic precursors (188 tryptic / 1155 non-tryptic), we quantified the intensity benefit at a depth of 5 transitions into the spectrum as observed in DI-MS (Fig. 2F). Signal intensity significantly improved using the optimized NCE compared to the standard NCE of 30% for 38.8% of tryptic peptides and 63.1% of non-tryptic peptides (p ≤ 5%, fold-change ≥ 1.5). Taken together, we show that optimizing NCE for optimal fragmentation efficiency increases both sensitivity and identification confidence for a large proportion of peptides and in particular non-tryptic peptides. The implemented DI-MS workflow is easily applicable for medium-to-large peptide sets and can be used to optimize virtually any MS parameter.

**Figure 2:**
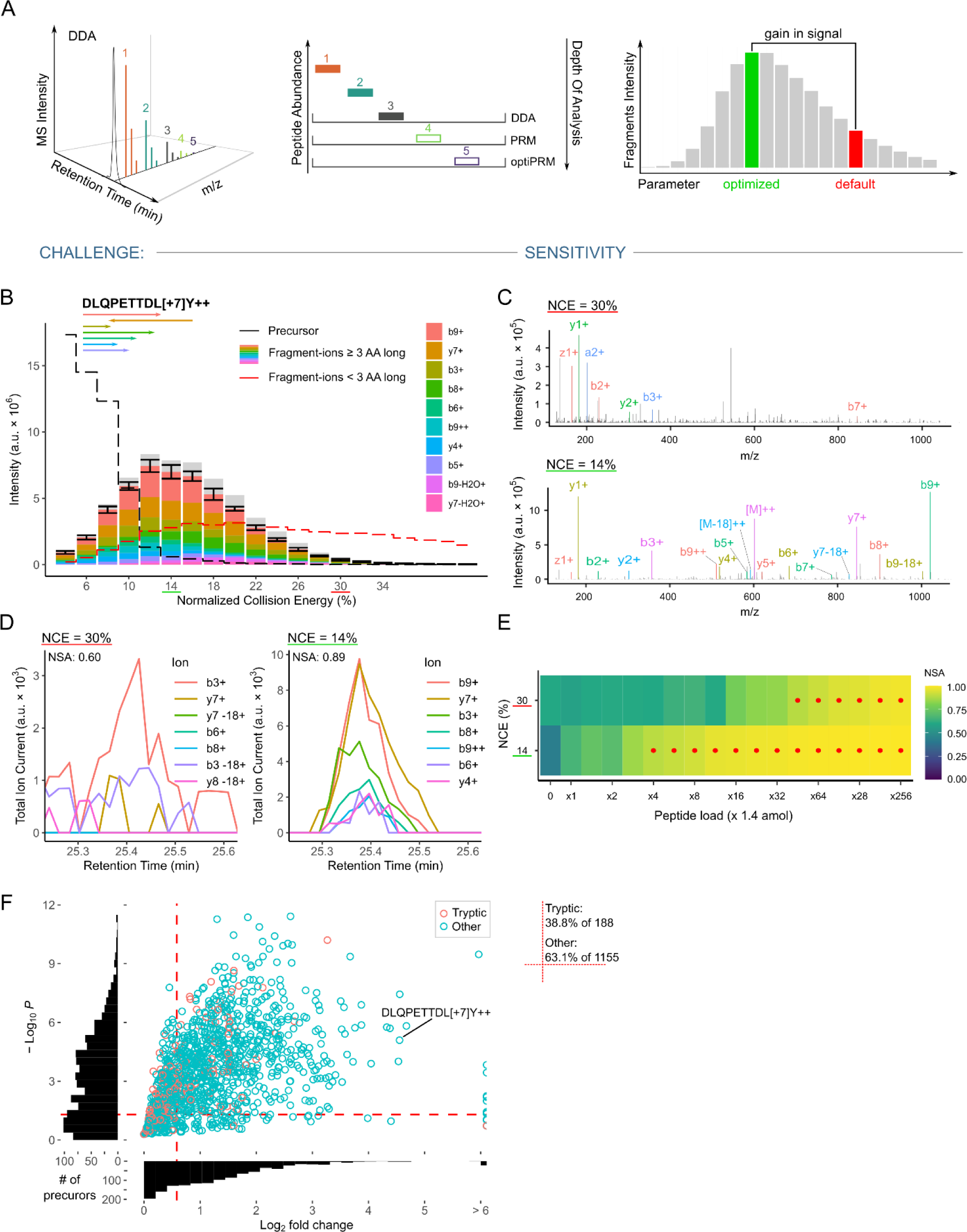
Collision energy optimization for target peptides. (A) Schematic overview of limitations of DDA and unoptimized PRM methods. Parameter optimization enables improved depth of analysis and is performed by systematically scanning parameter values and selecting the value with the best intensity response. (B) Data acquired for normalized collision energy (NCE) optimization via direct infusion illustrated for the peptide DLQPETTDL[+7]Y at precursor charge state +2. The arrows underneath the peptide sequence indicate b- and y-ions. For each NCE, four spectra were acquired and the acquired mean fragment intensities are shown. A set of dominant transitions of length ≥ 3 AA are shown in color. The standard deviation is given for the sum intensity of the indicated transitions. Sum intensity of precursor ions and transitions < 3 AA are shown as dashed black and red lines, respectively. The default NCE of 30% is marked in red and the optimized value for the peptide of 14% is marked in green. (C) Fragmentation (MS2) spectra at default and optimized NCE for the peptide in (B) acquired in an LC-MS PRM run. Both spectra were acquired close to the peak maximum (< 1 s apart). All transitions that could be matched with 7 ppm tolerance are annotated. (D) Extracted fragment ion chromatograms from a single LC-MS run alternating between the two NCE settings. 5.5 amol of the peptide in (B) was loaded with 50 ng HeLa tryptic digest to produce a signal close to the limit of detection. Transitions of length ≥ 3 AA are shown. (E) Normalized spectral contrast angle (NSA) results comparing acquired peaks to a reference library in a dilution series down to 1.4 amol of peptide in (B). A background matrix of 50 ng HeLa tryptic digest was used for all runs. High confidence detections with an NSA ≥ 0.85 are marked with a red dot. (F) Direct infusion NCE optimization result for all relevant precursors, quantified as the gain in signal intensity at a depth of 5 transitions into the spectrum. Compared is the signal at a standard NCE of 30% to the optimal value for each precursor. Histograms for the x- and y-axis show the overall distribution of points (peptides) in the plot. Peptides in the top-right corner, delimited with dashed red lines, show a signal gain ≥ 50% at a p-value < 0.05 (unpaired one-sided t-test). The data point corresponding to the peptide analyzed in (B) – (E) is indicated. The percentage of peptides appearing in the top-right section and an overall count are indicated to the right of the main panel.

### Optimized PRM parameters allow the detection of mutation-derived neoepitopes from limited input material

To test our optiPRM workflow, we applied it to detect mutation-derived neoepitopes from a patient-derived xenograft (PDX) pancreatic cancer cell line. Whole exome sequencing and RNA sequencing were carried out for the PDX cell line followed by mutation calling. Mutated protein sequences were generated and subjected to HLA binding prediction using NetMHCpan 4.1 (Fig. 3A). We curated a list of target peptides based on the following parameters: predicted binding to either HLA-A or HLA-B; exclusion of peptides where the corresponding wildtype peptide was also a predicted binder and the amino acid change occurred in an anchor position or outside the TCR recognition region; exclusion of peptides containing a cysteine residue. In total, we selected 55 peptides and synthesized their stable isotope labelled (SIL) variants for NCE optimization using DI-MS. The peptides were subsequently acquired using LC-MS in PRM mode for validation and to allow for optimal targeting. As a starting point, we performed optiPRM for samples originating from 1×10^8^ million cells, optionally treated with IFNγ (Fig 3B – E). Here, the high-resolution and high mass-accuracy acquisition on the Orbitrap MS with narrow precursor isolation (0.7 *m/z*) produces highly specific signals when fragment ion chromatograms are extracted with 7 ppm tolerance. To evaluate peptide detection, we found the most informative value to be the NSA of the detected transitions compared to the SIL reference and applied a stringent cutoff of NSA ≥ 0.85. Validity of the detections was additionally ensured by manual peak curation requiring a minimum of 5 detected transitions and an LC retention time matching the SIL reference. A panel of NSA results for the top 30 binders (based on %EL rank) targeted in this experiment is shown in Fig. 3B with red dots indicating confident identifications with an NSA ≥ 0.85. We detected the peptide RIAESLPVV in both conditions (± IFNγ) and biological replicates (Fig. 3B – D). The peptide SRFTGATII was found at the detection limit, appearing only in one biological replicate (Fig. 3E). To evaluate the sensitivity of our optiPRM assay, in an independent experiment the assay was scaled down to a minimum of 2.5×10^6^ cells input material. Again, we detected the peptide RIAESLPVV in all samples and technical MS replicates. The second detected peptide SRFTGATII was not detected using lower sample amounts.

**Figure 3.**
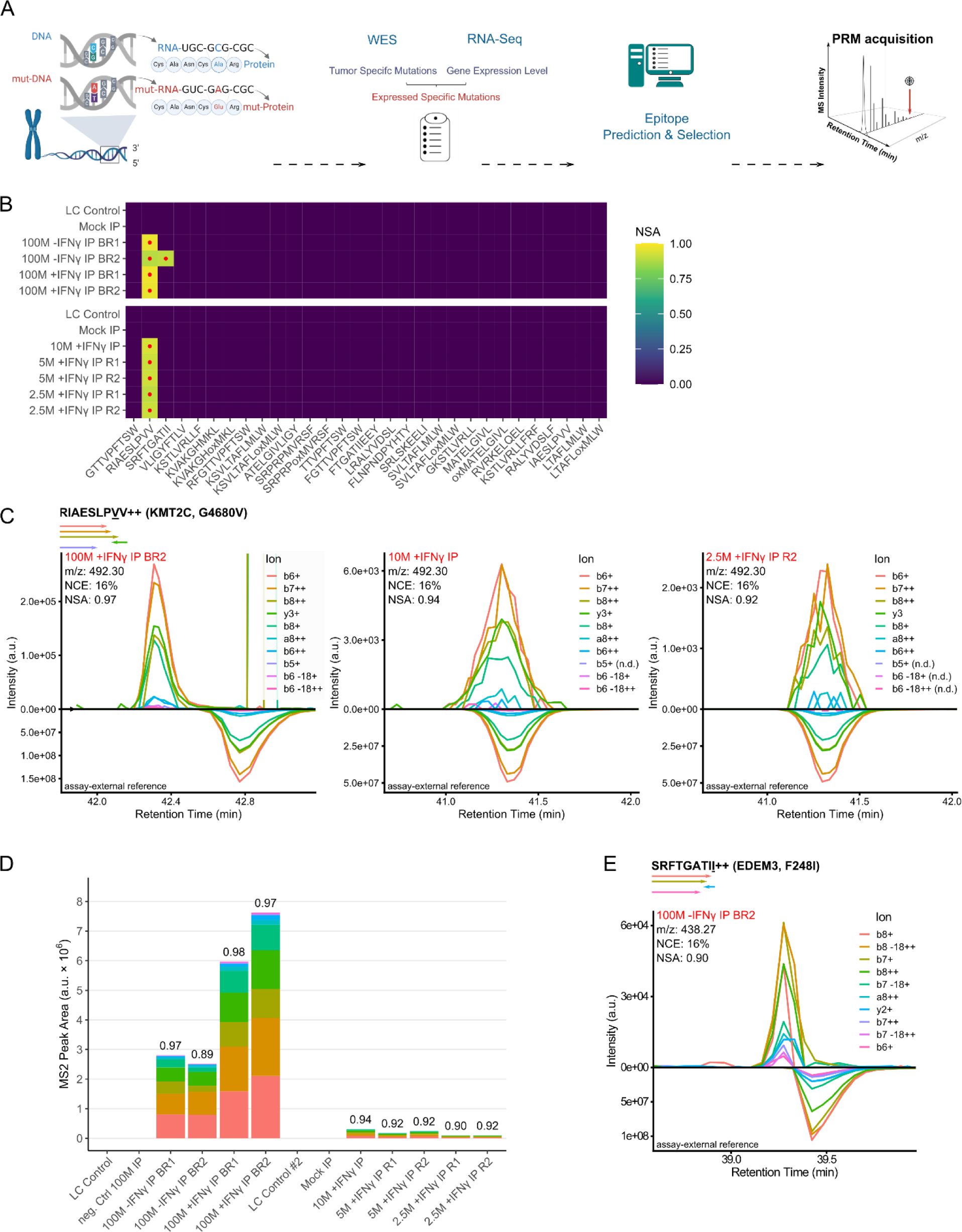
Sensitivity of neoepitope detection in a primary human patient-derived xenograft (PDX) cell line. (A) Schematic overview of the targeted workflow for neoepitope detection in tumor samples. An example of a point mutation (SNP) is depicted in the left panel. PDX cell lines were submitted to whole exome sequencing (WES) and RNA sequencing. Identified mutations with sufficient mutant allele frequency and expression level were submitted to epitope prediction. Selected epitope candidates were targeted in LC-MS using PRM. (B) Cell plot showing an overview of the targeted LC-MS (PRM) results. Peptide sequences are shown for the top 30 candidates sorted by predicted netMHCpan 4.1 eluted ligand percentile rank (lowest to highest). The normalized spectral contrast angle (NSA) compares acquired fragment ion peaks in a sample to the reference library. High confidence detections with an NSA ≥ 0.85 are marked with a red dot. Negative controls LC Control (PRTC standards on BSA background) and Mock IP (IP replicate without cell input) were run immediately prior to the samples. An initial series with 100M cell load immunoprecipitations (IPs) injected in full and a successive series going down to 2.5M cell load are depicted. BR, biological replicate; R, technical replicate. (C) Extracted fragment ion chromatograms (XICs) for peptide RIAESLPVV resulting from decreasing IP cell load. XICs compare peaks from sample runs (up) and the corresponding stable isotope labelled (SIL) synthetic reference peaks (down), acquired separately. Reference peaks are aligned on the RT axis by linear interpolation between two adjacent reference peptides (PRTC Standard) acquired in both runs. Plausible detections are scored by NSA ≥ 0.85. The name of the mutated gene, the wild-type/mutated amino acid and its position are indicated in the bracket following the peptide sequence. The arrows underneath the peptide sequence correspond to the detected b- or y-ions with the same colors as in the chromatograms. (D) Relative quantification for the peptide RIAESLPVV in all replicates. Total bar height corresponds to the sum intensities of fragment ions indicated with the same colors as in the chromatograms. (E) XIC for detection of peptide SRFTGATII in a single replicate. Illustration as in (C).

Comparison of the XICs for RIAESLPVV shows a gradual loss of transition detections (marked n.d. (not detected) in Fig. 3C) with lower cell load. At 2.5×10^6^ cells, 3 of the lowest intensity transitions were not detected anymore, while the NSA remained high at 0.92, indicating the high quantitative similarity of the remaining transition signals to the synthetic reference. In addition, the XIC only shows signal clearly attributable to the detected peptide, even though it is close to the limit of detection. SIL peptides are commonly added to the sample as an internal control to compare the sample signal to the synthetic reference. However, we observed light contamination of SIL peptides ^52^. This is a major concern when targeting lower abundance peptides, as it increases the risk of false detection (Fig. S2A). Consequently, we decided against adding SIL analogs of targeted neoepitopes to our samples. During method establishment and testing using the PDX cell line model, we noticed several further potential pitfalls that might lead to false-positive identifications. To avoid contamination, we never used unlabeled synthetic target peptides in our LC-MS system. It is nevertheless a requirement to wash the system after exposing it even to labelled peptides and diligently test for their presence before analyzing an IP sample (Fig. S2A). Contamination can also stem from similar IP samples, which can introduce target peptides into the system (Fig. S2B). To prevent such cases of false positive detection, we always performed negative control “LC Control” runs (BSA tryptic digest with PRTC standard) before the actual sample runs in this study. To ensure similar comparison integrity as with internal control, we mirrored reference XICs below the sample XICs and used an extra alignment step on the RT axis. This step corrects for common retention time shifts that appear during separate LC runs ^53^. The reference peak is aligned by linear interpolation between two adjacent reference peptides (PRTC Standard) acquired in both runs. In Fig. 3C, this results in completely aligned XICs for the center and right panels (reference shifted −0.5 min). In contrast, the left panel with the strongest target signal shows a deviation between sample and reference peaks (reference shifted −1.2 min). While the alignment is beneficial in all three cases, this indicates limitations to the technique, likely due to non-linear local RT shifts ^53^. In addition to fragment ion chromatograms, we extracted precursor ion chromatograms from full MS scans, but found that they did not reach the same sensitivity (Fig. S3A). Label-free quantification of fragment ions of the peptide RIAESLPVV revealed more than 10-fold signal reduction when going from 1×10^8^ to 1×10^7^ load (Fig. 3D). Expectedly, in samples treated with IFNγ, signal is increased compared to the untreated control samples, which can be attributed to an induced increase in HLA expression. Taken together, our optiPRM workflow can be used to detect low abundant, mutation-derived neoepitopes from limited sample amounts. Notably, we were able to detect one neoepitope not only in samples from the commonly used input amount of 1×10^8^ cells but also in samples corresponding to as little as 2.5×10^6^ cells.

### Detection of mutation-derived neoepitopes in tumor biopsies

After proving the enhanced sensitivity of our optiPRM workflow we next utilized it to analyze tumor biopsies of varying size from five patients (Table S1). We performed whole exome sequencing and RNA sequencing for all patients to identify mutations derived from SNVs, InDels and fusions. The resulting peptide sequences were *in silico* queried with NetMHCpan 4.1 for potential HLA class I binders. Candidate peptides selected for DI-MS parameter optimization were curated using the previously described criteria. The first sample analyzed had a tumor cell content > 90%, weak to moderate HLA class I expression and originated from a patient (female; 18 years) with osteosarcoma and lung metastasis. In total, we selected 43 candidate peptides to be targeted by our optiPRM assay leading to the detection of two mutation-derived neoepitopes: RYIGDAHTF and RYIGDAHTFAL, both originating from an SNV in *ARHGAP35* (p.R211G)(Fig. 4A & D). Both peptides are predicted to strongly bind to HLA-A*23:01 with a %EL rank of 0.002 and 0.193, respectively. The second analyzed sample had a tumor cell content of 50 – 80% with moderate to strong HLA class I expression and originated from a patient (male; 55 years) with a small intestine carcinoma. From 49 selected candidate peptides, one (APRQPLSSI) was detected in the tumor biopsy (Fig. 4B & E). The peptide derives from an SNV in *RNF111* (p.S211R) and is predicted to strongly bind to HLA-B*56:01 (%EL rank: 0.072). Of note, the peptide was detected in two injections from just 35 mg of tissue as input material, further highlighting the ultra-high sensitivity of our assay. The third sample had a tumor cell content of 10 – 20%, weak to moderate HLA class I expression, and originated from a patient (female; 51 years) with liposarcoma. The mutational landscape of this patient was characterized by a high number of fusion events involving chromosome 12, a condition found to be common in liposarcoma ^54^. From 66 selected candidates, a subset of the corresponding synthetic surrogate peptides turned out to elute at exceptionally late retention times (Fig. S4A) indicating a strong hydrophobic character, which typically is not compatible with elution using only 28% ACN from SepPak cartridges. To account for these peptides, we tested a recently published alternative solid-phase extraction technique with restricted access material (RAM). RAM allows elution with higher ACN percentages and therefore better performance with late eluting peptides ^25^. Using a cell line model, we compared RAM to our standard C18 desalting and indeed observed improved detection of late eluting hydrophobic peptides (Fig. S4B – C). In contrast to the previously published protocol, we found that loading the sample in acidic conditions allowed to identify the highest number of peptides. In particular, it allowed for the detection of more peptides with a low isoelectric point compared to the standard loading in neutral conditions (Fig. S4D). Such peptides are expected to be negatively charged at pH 7, which may prevent binding to the RAM material, which we suspect to be also negatively charged at neutral pH. Operating RAM in acidic conditions avoids this limitation. Analysis of the tumor sample with RAM resulted in the detection of two of the 66 candidates tested (Fig. 4C). The first peptide, ARFMSPMVF, originates from a fusion involving *TSPAN8* and *CPM* (Fig. 4F). The peptide is predicted to strongly bind to HLA-B*27:05 (%EL rank: 0.015) and HLA-C*07:02 (%EL rank: 0.026). The second peptide, RRIRASQLLLH, originates from a fusion involving *PAPOLA* and *MGAT4C* and strongly binds to HLA-B*27:05 (%EL rank: 0.29; Fig. 4G). For both peptides, the first replicate provided better signal intensity and NSA. In contrast, the second sample, at a lower intensity, indicates exact coelution with the reference, similar to what was observed for the PDX example. Again, precursor ion chromatograms did not reach the same sensitivity for all but one precursor (Fig. S3B). For patients 4 and 5 we did not detect any of the *a priori* selected neoepitope candidates using the optiPRM workflow. Taken together, we identified five distinct mutation-derived neoepitopes from three of the five clinical samples tested using our optiPRM assay (details summarized in Table S3).

**Figure 4.**
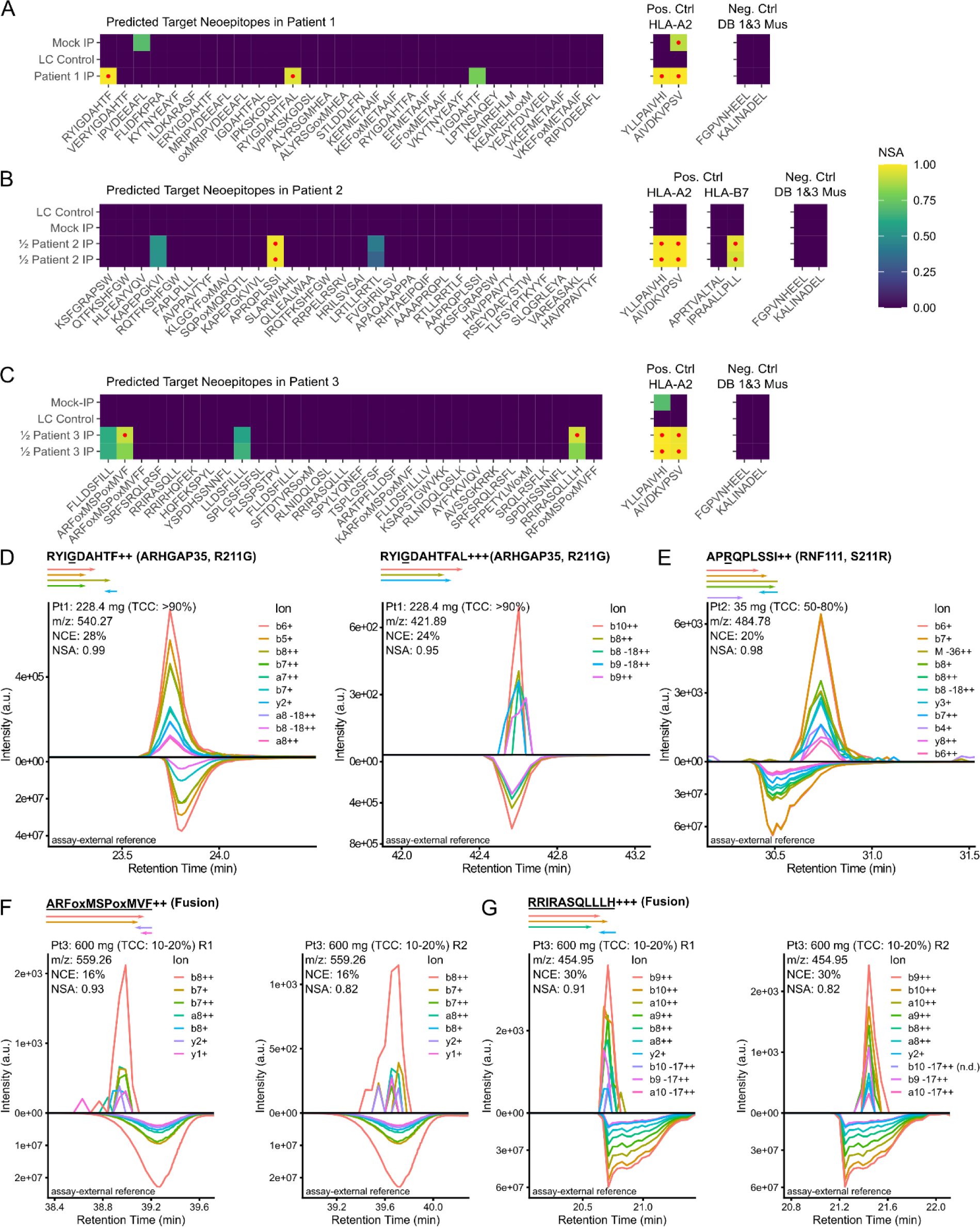
Targeted LC-MS analysis of three patient tumor biopsies. Presentation of the results according to the same metrics and color code conventions as presented in Fig. 3. (A) – (C) Cell plots summarizing the results for three patient samples. Peptide sequences are shown for the top 30 candidate epitopes sorted by predicted netMHCpan 4.1 eluted ligand percentile rank (lowest to highest). In addition, a selection of common binders for the targeted HLA supertypes is included as positive controls and a set of mouse epitopes as negative controls. High confidence detections with an NSA ≥ 0.85 are marked with a red dot. Negative controls LC Control (PRTC standards on BSA background) and Mock IP (IP replicate without cell input) were run immediately prior to the samples. IPs from patients 2 and 3 were injected in two equal-sized technical replicates. (D) – (G) XICs for the peptides detected in (A) – (C). The sequence alterations compared to the wild-type are underlined in each investigated peptide. For SNPs, the gene name is followed by the mutated amino acid position. Peptides with frameshift sequences derived from gene fusions are marked “Fusion”. Patient samples are indicated as Pt1, Pt2, or Pt3 for Patients 1, 2, and 3 with tumor sample weight and sample tumor cell content (TCC).

### Mutation-derived neoepitopes are recognized by autologous CD8^+^ T cells

To assess the existence of CD8^+^ T cells recognizing the mutation-derived neoepitopes, we performed MHC multimer staining of autologous peripheral blood mononuclear cells (PMBCs) directly *ex vivo* and after 10 days of *in vitro* stimulation with the cognate peptide. For the peptides RYIGDAHTF and RYIGDAHTFAL detected in patient 1 (data not shown) as well as for the peptide APRQPLSSI detected in patient 2 (Fig. 5A), we could not detect CD8^+^ T cells recognizing the corresponding MHC multimers. A possible explanation for the lack of specific CD8^+^ T cells could be the high similarity of these neoepitopes with their corresponding wild-type peptide. We assessed this exemplarily for APRQPLSSI bound to HLA-B*56:01 by *in silico* modeling. Comparing the models for APRQPLSSI and the corresponding wild-type peptide APSQPLSSI reveals that the arginine side chain introduced by the mutation is buried in the peptide binding groove of the HLA molecule and therefore likely not accessible for a TCR (Fig. 5B). In contrast, the molecular surface of the HLA:peptide complex is very similar and thus probably indistinguishable between wild-type and mutant peptide and consequently let to a thymic depletion of TCRs recognizing these epitopes.. For the third patient, peptide-specific autologous CD8^+^ T cells were detected for the fusion-derived neoepitope ARFMSPMVF in the context of HLA-B*27:05 (Fig. 5C). The neoepitope was also predicted to bind to HLA-C*07:02, however we did not detect CD8^+^ T cells recognizing this HLA:peptide combination or the second fusion-derived neoepitope RRIRASQLLH detected by mass spectrometry. Of note, the CD8^+^ T cell population recognizing ARFMSPMVF was only detectable after *in vitro* stimulation but not directly *ex vivo*.

**Figure 5.**
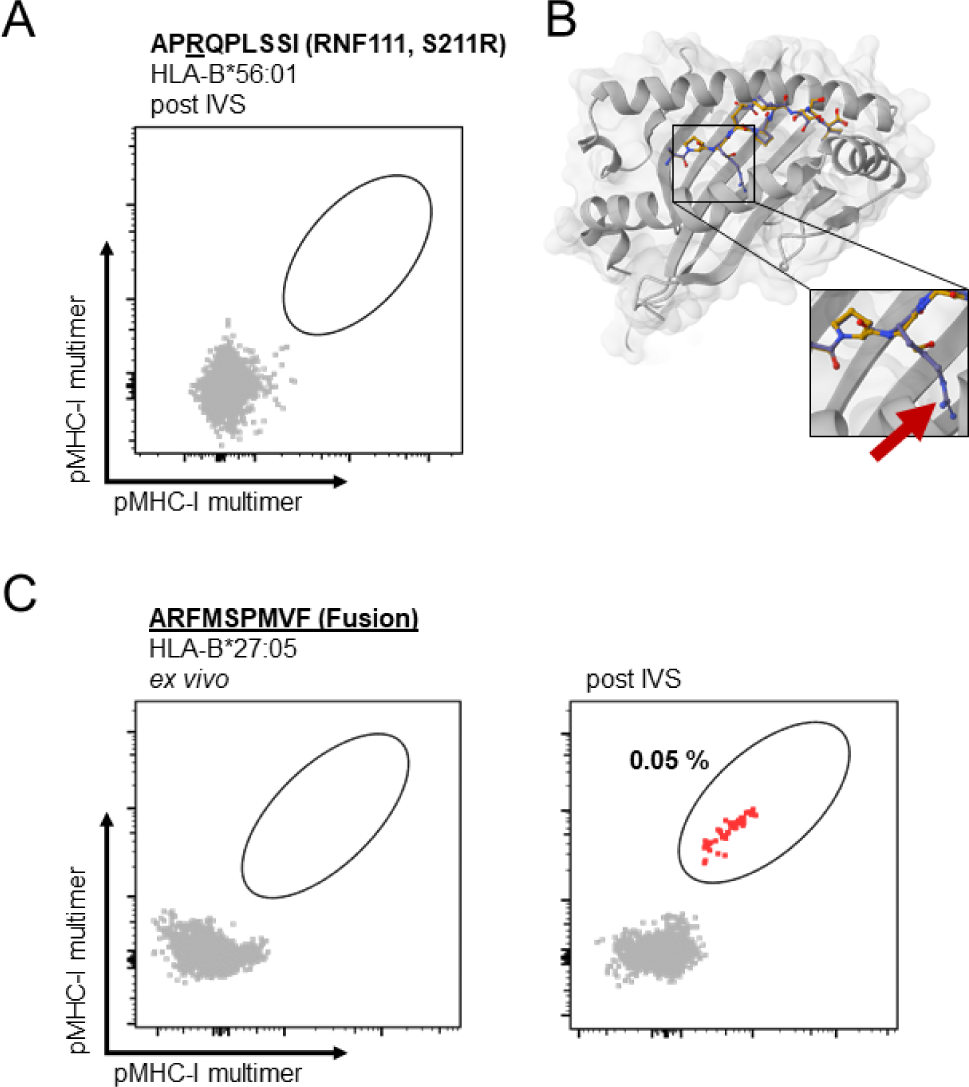
HLA:peptide multimer staining and structural modelling of mutation-derived neoepitopes. (A) HLA:peptide multimer staining results of patient-autologous PBMC recognition of APRQPLSSI bound to HLA-B*56:01 post peptide-specific *in vitro* stimulation (IVS). (B) 3D structural model created with PANDORA comparing binding of the wildtype peptide (APSQPLSSI; yellow) and the neoepitope (APRQPLSSI; purple). The red arrow indicates the predicted position of the R residue inside the HLA binding groove. (C) HLA:peptide multimer staining results of patient-autologous PBMC recognition of ARFMSPMVF bound to HLA-B*27:05 *ex vivo* and post peptide-specific IVS.

## CONCLUSIONS

Identification of targetable epitopes is the most crucial prerequisite for the development of safe and efficient immunotherapies such as therapeutic cancer vaccines or TCR-transgenic immune cell products. Mass spectrometry-based immunopeptidomics is the only method available to provide direct proof of actual epitope presentation at the surface of tumor cells. Here, untargeted MS acquisition schemes such as DDA or DIA are the predominant methods used for the identification of targetable epitopes, while targeted methods are mainly used for confirmation purposes ^2,55^. However, DDA and, to a lesser extent, DIA lack the unrivaled sensitivity achievable by targeted methods ^56^ and which is needed for the detection of particularly low abundant mutation-derived neoepitopes or viral epitopes ^18,20^. To this end, we developed optiPRM, a targeted MS workflow utilizing PRM on Orbitrap instrumentation, to detect HLA class I-presented peptides from limited sample amounts. The ultra-high sensitivity of the optiPRM approach is achieved by systematic optimization of acquisition parameters on a per peptide basis – a process which is so far unattainable for untargeted methods. The use of synthetic peptides in advance allows for tuning of MS acquisition parameters but also serves as empirical reference data that can be directly compared to the acquired signal to ensure that detections can be confidently asserted even at very low intensity levels. While using SIL peptides as an internal standard in proteomics is common for high-confidence validation and quantification, we have previously found this to be affected by contamination with unlabeled target peptides and therefore advise against this method for the confirmation of especially low-intensity targets such as mutation-derived neoepitopes ^52^. Consequently, we have opted for comparing assay-external reference chromatograms acquired in separate runs and aligned on the retention time axis based on adjacent standard peptides. This offers many of the same benefits, allows exclusive use of instrument acquisition time on epitope candidates and was very much in agreement for most of our detections. However, it can fall short in some cases, such as for the neoepitope detections from PDX samples with high input amounts. There, we observed a strong general retention time shift. While this was corrected for, it additionally seemed to induce further non-linear retention time distortions and peptide elution order inversions. These challenges for our retention time alignment are consistent with what has been described in the literature ^53^. Here, further refinement through an extended set of standard peptides or non-linear interpolation may be feasible. In this study, we show that optimizing normalized collision energy used for peptide fragmentation significantly increases sensitivity, especially for non-tryptic peptides such as HLA class I-presented peptides. We show that detecting mutation-derived neoepitopes is possible from as little as 2.5×10^6^ cells or 35 mg tumor tissue as input, which is roughly one to two orders of magnitude less than commonly used in immunopeptidomics protocols ^50^. While we focused on optimizing normalized collision energy in this study, our direct injection approach is applicable to virtually any MS parameter and could, for example, be used to optimize ion optics transmission parameters such as radio frequency (RF) lens voltages or FAIMS compensation voltages for maximum transmission of peptides of interest. These are independent from optimization of collision energies and can further enhance sensitivity. In total, we targeted 274 peptides in 5 different patient samples using the optiPRM method and were able to identify 5 distinct mutation-derived neoepitopes, while we did not detect mutation-derived neoepitopes in two other patients analyzed in this study. Comparing this with the estimation of one detectable neoepitope per 1.8×10^8^ non-synonymous mutations ^57^ highlights the success rate of our approach. However, it also exemplifies some limitations of mass spectrometry-based immunopeptidomics in general and targeted approaches in particular. In general, mass spectrometry might not detect some peptides due to their chemical properties and/or their low abundance below the detection limits for current instrumentations. Additionally, *a priori* candidate selection introduces a bias in targeted immunopeptidomics approaches. It is especially true when considering *e.g.* HLA binding predictions of poorly investigated HLA alleles as selection criteria, which can exclude actually presented peptides. Conversely, targeted methods offer superior performance given a sufficiently small set of targets ^58^. Indeed, most of our neoepitope detections showed little to no MS1 signal, likely making them undetectable by untargeted approaches. While PRM methods can target up to several hundreds of peptides ^59^, we stayed well below such counts (*i.e.* ≤ 66) to ensure few tradeoffs are made with regards to sensitivity. For the manual selection of candidates, we mainly focused on *in silico* HLA binding predictions but also considered RNA expression data and dissimilarity to the corresponding wildtype peptide where applicable. In the future, the success rate of targeted immunopeptidomics workflows may be further improved as the predictive power of such prioritization improves due to the continuing advancements in understanding antigen processing and presentation ^46,60–62^. Here, especially the growing amount of available immunopeptidomics data for rare and thus poorly investigated HLA alleles might boost the predictive power. A reason why studies solely relying on therapeutics against *in silico* <u>predicted</u> neoepitopes ^63–65^ have been only partly successful so far is the possibility that none of the selected neoepitopes are presented at the surface of the tumor cells. While mass spectrometry-based immunopeptidomics is currently the only available technique to provide direct proof of actual neoepitope presentation by HLA molecules, these neoepitopes must be recognizable by CD8^+^ T cells to trigger an immune attack leading to tumor elimination. To evaluate this, we screened patient-autologous PBMCs by MHC multimer staining for the HLA:peptide combinations detected. Using in-house produced soluble HLA:peptide complexes, we found T cell populations for only one of the neoepitopes, which was derived from a gene fusion and the corresponding frameshift, potentially showing very little “similarity-to-self”. Interestingly, this population was only detectable following *in vitro* T cell expansion. First, this exemplifies the clinical potential of the detected neoepitope as this population is expandable, indicating that the corresponding patient may benefit from a cancer vaccine targeting this peptide. Second, it highlights the merit of our targeted immunopeptidomics workflow as this neoepitope might have been missed in assays omitting the challenging procedure of T cell expansion. In the case of APRQPLSSI for which we did not detect specific T cell responses, this can be explained by *in silico* modeling of the HLA:peptide complex, revealing that the side chain of the mutated amino acid is placed deep inside the binding pocket and therefore likely does not affect TCR binding. Again, improvements in “similarity-to-self” prediction and incorporation of HLA:peptide modeling should enable *a priori* exclusion of such candidates with low likelihood to be recognized by T cells, especially for neoepitopes derived from single amino acid changes. Moreover, patient T cells against mutation-derived neoepitopes might be non-functional ^66^ and identification of reactive T cell clones could benefit by screening autologous healthy donors.

Cancer neoepitope identification pipelines are becoming a major endeavor in recent years, and progress is rapid on all fronts such as *in silico* prediction of epitope HLA-presentation and high-throughput scanning for T-cell recognition. However, MS-based immunopeptidomics remains the sole approach that provides the ultimate proof of neoepitope presentation at the surface of tumor cells. Our optiPRM method empowers the detection of neoepitopes with ultra-high sensitivity and from limited amounts of clinical samples. In the future, these validated neoepitopes might serve as starting point for the successful development of personalized cancer therapies such as vaccines or transgenic TCR T cell products.

## ACKNOWLEDGEMENTS

We are thankful for the excellent technical assistance of Rebecca Köhler (IPs, AB-Beads, gels), Sophia Föhr (LC-MS), Claudia Luckner-Minden, Iris Kaiser, Annette Köster, Selina Börsig (biobanking, sample processing), Rosa Eurich (immunohistochemistry) and Aaron Rodriguez Ehrenfried (TIPC expansion). We would like to acknowledge the excellent services and support provided by the DKFZ core facilities for genomics & proteomics and flow cytometry.

## AUTHOR CONTRIBUTIONS

Conceptualization: ABR; Methodology: MSa, JDF, MM, FM, IP; Investigation: MSa, JDF, JPB, MM, YL, KL, CL, MV; Formal analysis: MSa, JDF, JPB; Validation: MSa, JDF, JPB, MM, FM, IP, ABR; Software: JDF, PC, YL, MV; Visualization: MSa, JDF, JPB, MM; Data curation: JDF; Resources: SF, MSch, RO, MP, DJ, IZ, ABR; Funding acquisition: SF, MSch, RO, MP, DJ, ABR; Project administration: IZ, ABR; Supervision: ABR; Writing - Original Draft: MSa, JDF, JBP, ABR; Writing - Review & Editing: MSa, JDF, JBP, ABR. All authors have read and agreed to the manuscript.

