## Supplementary material for "optiPRM: A targeted immunopeptidomics LC-MS workflow with ultra-high sensitivity for the detection of mutation-derived tumor neoepitopes from limited input material": Supplemenal Figures

**SUPPLEMENTARY FIGURES**

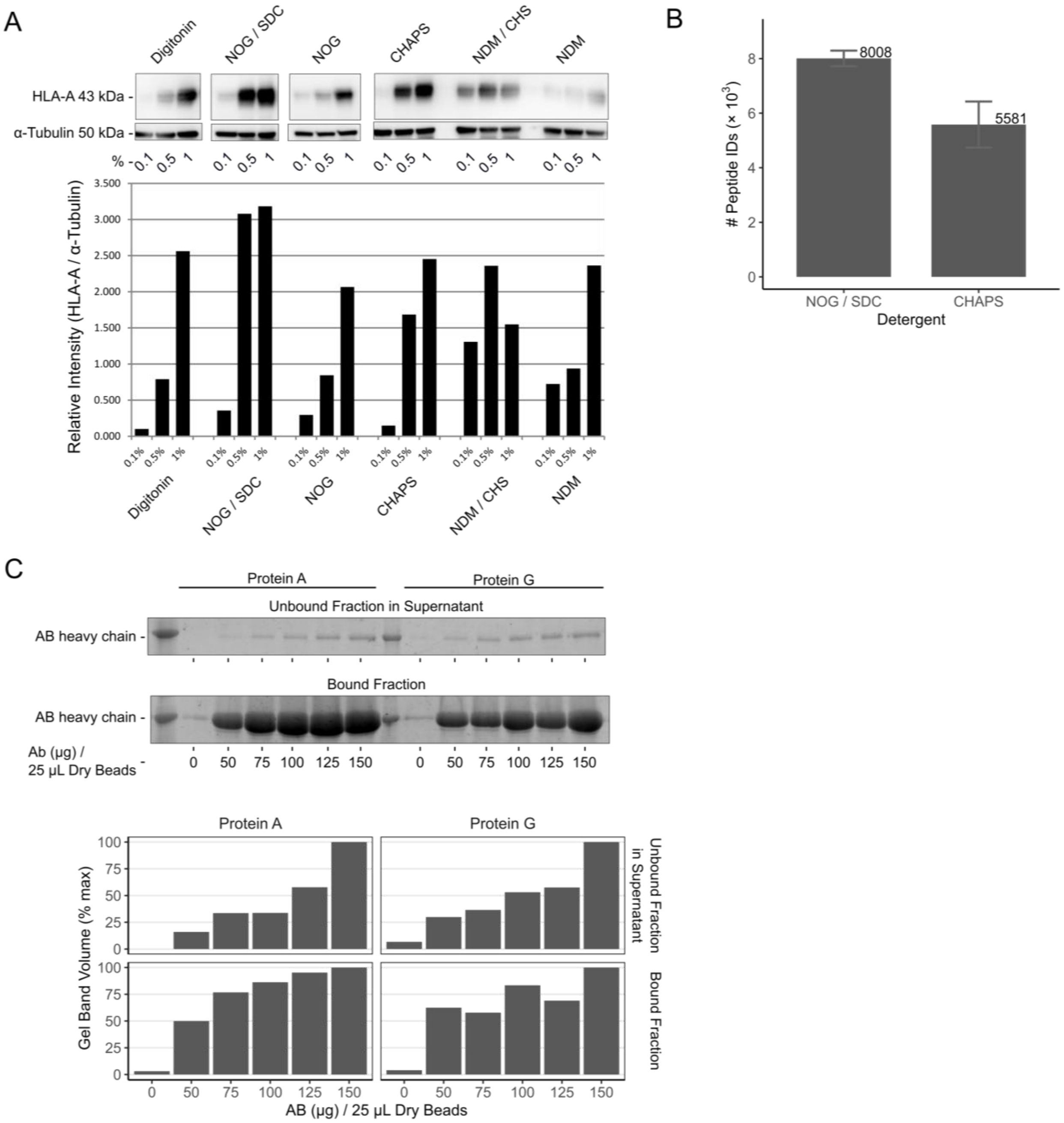


**Figure S1.** Revisiting key parameters of the HLA:peptide immunoprecipitation

(A) Anti HLA-A immunoblot after cell lysis in buffers with different non-polymeric detergents (non-ionic: n-octyl-β-D glucopyranoside (NOG), n-dodecyl β-D-maltoside (NDM) and cholesteryl hemisuccinate (CHS); biological: digitonin and sodium deoxycholate (SDC); zwitterionic surfactant: 3-[(3-cholamidopropyl)dimethylammonio]-1-propanesulfonate (CHAPS)) at three different volume percentages. Tubulin: loading control.

(B) Number of unique peptides identified by LC-MS in DDA mode from anti-panHLA IPs carried out at 1% CHAPS versus 1% NOG in combination with 0.25% SDC. In each condition, the results represent three independent replicates, each run in two technical replicates. The average number of unique peptides identified in each condition is indicated. The error bars represent the standard error of the mean (+/-).

(C) SDS-PAGE gel (stained with Oriole Fluorescent Gel Stain®, BioRad), from an experiment testing the saturation of 25µL dry protein A or protein G beads with the indicated increasing amount of anti-panHLA Ab (clone W6/32). The upper gel shows the Ab band leftover in the cell lysate after coupling with beads (Supernatant Post-Coupling). The lower gel shows the bound fraction (Ab Post-Coupling bound to protein G beads) eluted by boiling the beads. For relative quantification, band intensities were compared to corresponding samples with 150 µg AB/25µl dry beads.

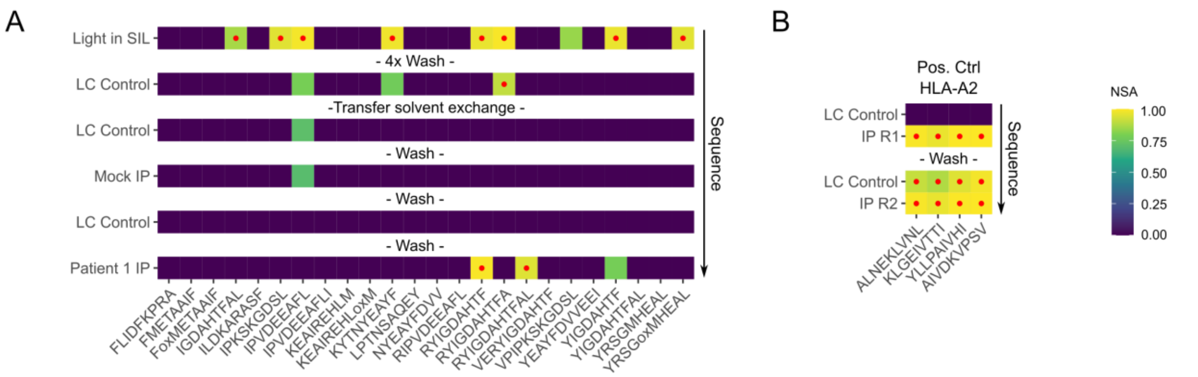


**Figure S2.** False detection concerns due to trace-level carry-over on the LC system

NSA (normalized spectral angle) cell plot for target (light) peptide detections. High confidence detections with an NSA ≥ 0.85 are marked with a red dot.

(A) Light peptide detections in successive LC-MS runs after synthetic SIL peptide acquisition. Sequential acquisition from top to bottom. The initial run “Light in SIL” depicts light peptide detections due to synthetic SIL peptide reference acquisition. LC control (PRTC standards on BSA background) was run to monitor persistent contamination after successive washes leading to eventual clearance for sample IP acquisition.

(B) Carryover of the positive control signal from IP replicate 1 to the LC control run prior to replicate 2.

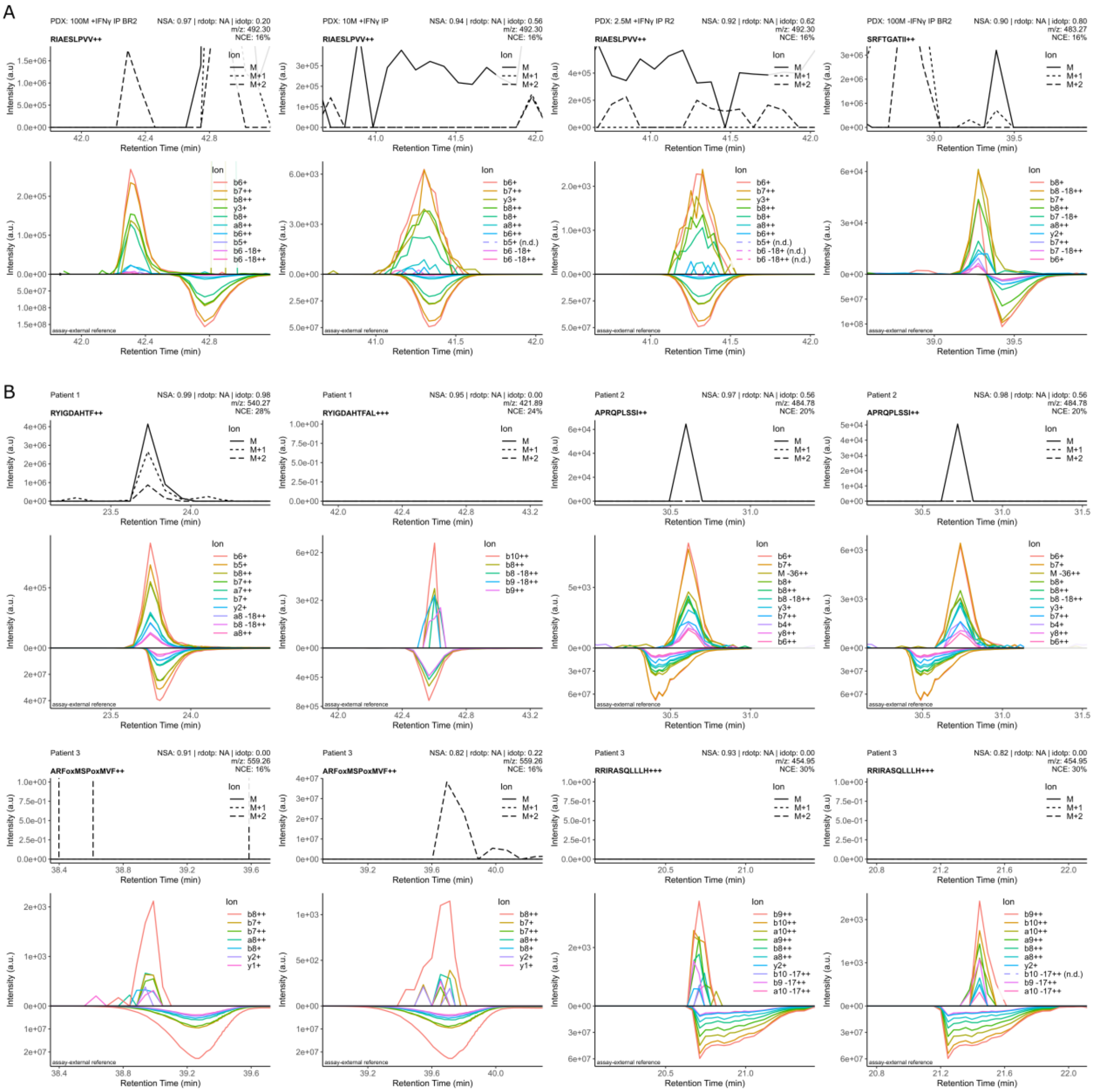


**Figure S3.** Full MS1 precursor chromatograms for all detections

(A) and (B) Top panel shows precursor (full MS1 based) XICs belonging to the epitope chromatograms presented in Figure 1 and 2. Annotated skyline metadata: NSA (referred to in skyline as dotp), rdotp is not used, idotp is NSA for precursor XIC vs. theoretical analyte isotopic envelope.

(A) XICs for detections in PDX

(B) XICs for detections in patient samples

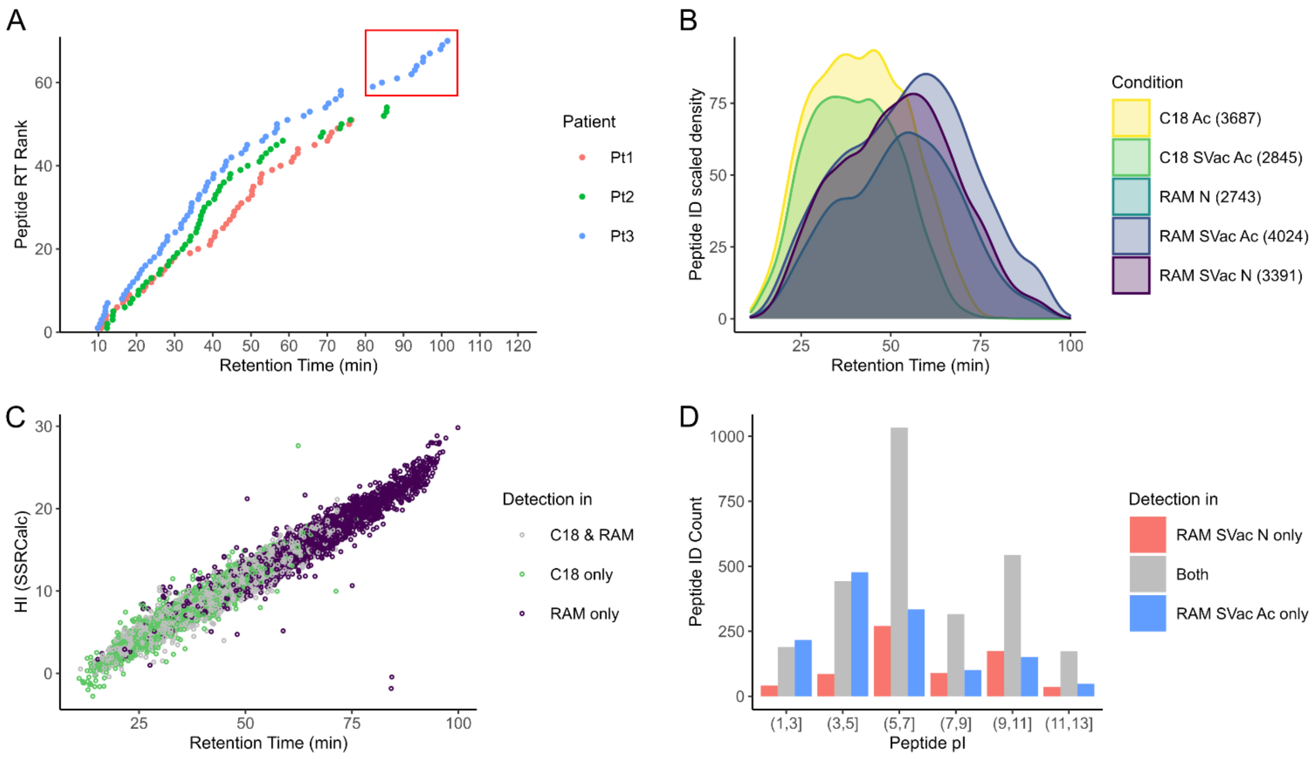


**Figure S4.** Analysis of peptide purification with restricted access material (RAM)

A) Target peptide retention times for Patients 1, 2, and 3 as determined by the acquisition of synthetic SIL variants. A substantial set of late eluting peptides for Patient 3 is marked with a red box.

B) Untargeted LC-MS peptide identifications. Scaled ID density as a function of retention time. C18 (SepPak) and RAM, with or without Speed Vac (SVac) drying before IP, loading in neutral (N) or acidic (Ac) conditions. The numbers in the brackets indicate the number of identified peptides at 1% FDR.

C) Hydrophobicity index (HI) calculated by Sequence-Specific Retention Calculator (SSRCalc) plotted versus RT for all identified peptides resulting from C18 (SepPak) and/or RAM workflows.

D) Peptide identification counts for RAM loading in neutral (N) and/or acidic (Ac) conditions. Peptides are grouped by isoelectric point (pI) ranges to reveal the divergence in performance at low pI.

**SUPPLEMENTARY TABLES**

**Table S1.** Patient information including ethics vote information, malignancy and sample properties

**<Table S1 is a separate excel file: Table S1.xlsx>**

**Table S2.** MS run specifications for the relevant LC-MS assays presented in this study

**<Table S2 is a separate excel file: Table S3.xlsx>**

**Table S3.** Epitope specifications

**<Table S3 is a separate excel file: Table S3.xlsx>**
